# Multiple nutritional and gut microbial factors associated with allergic rhinitis: the Hitachi Health Study

**DOI:** 10.1101/2021.10.11.463208

**Authors:** Yukari Sahoyama, Fumiaki Hamazato, Manabu Shiozawa, Tohru Nakagawa, Wataru Suda, Yusuke Ogata, Tsuyoshi Hachiya, Eiryo Kawakami, Masahira Hattori

## Abstract

Several studies suggest the involvement of dietary habits and gut microbiome in allergic diseases. However, little is known about the nutritional and gut microbial factors associated with the risk of allergic rhinitis (AR). We recruited 186 apparently healthy workers with symptoms of AR and 106 controls at the Hitachi Health Care Center, Japan. The habitual consumption of 42 selected nutrients were examined using the brief-type self-administered diet history questionnaire. Faecal samples were collected and subjected to amplicon sequencing of the 16S ribosomal RNA gene hypervariable regions. Association analysis revealed that four nutrients (retinol, vitamin A, cryptoxanthin, and copper) were negatively associated with AR. Among 40 genera examined, relative abundance of *Prevotella* and *Escherichia* were associated with AR. Furthermore, significant statistical interactions were observed between retinol and *Prevotella*. The age- and sex-adjusted odds of AR were 25-fold lower in subjects with high retinol intake and high *Prevotella* abundance compared to subjects with low retinol intake and low *Prevotella* abundance. Our data provide insights into complex interplay between dietary nutrients, gut microbiome, and the development of AR.

## Introduction

Allergic rhinitis (AR) is a symptomatic disorder of the nose caused by immunoglobulin E (IgE)-mediated reactions against inhaled allergens^1,2^. The classical symptoms of AR are nasal itching, sneezing, rhinorrhoea, and nasal congestion, and ocular symptoms are also frequent^3^. Over 500 million people worldwide are suffering from AR^1^. The burden of disease related to AR is substantial due to reduced patients’ quality of life (QoL), lower performance at work and school, poorer sleep quality and cognitive function, and increased irritability and fatigue^3,4^. Moreover, AR is a risk factor for asthma^3^. Despite recognition of AR as a global health problem, its aetiological risk factors have yet to be fully understood^1^.

The prevalence of allergic diseases, including AR and asthma, has increased in many countries, and changes in dietary habits may be one of the factors responsible for this trend^5^. Diet is an important source of nutrients that may modulate the risk of allergic diseases^6^. For instance, systemic oxidative stress enhances inflammatory responses relevant to allergy; and therefore, antioxidants may play a key role in the prevention of allergic diseases^7^. Furthermore, epidemiological studies have suggested that fruits and vegetables that contain several antioxidants protect against allergic diseases^8^. Therefore, researchers have intensively investigated the nutrients exhibiting a protective (or adverse) effect on the development of allergic diseases, such as asthma^5–8^. However, little is known about the relationship between nutrients and the risk of AR.

Recent studies have suggested that gut microbiota may be involved in the pathogenesis of allergic diseases. The gut microbiota is mainly composed of bacteria, and its composition is highly dynamic and influenced by dietary habits of their host^9^. The gut microbiota produces bioactive metabolites from dietary materials^10^. These metabolites influence host immune responses through the interplay with regulatory T (Treg) and dendritic cells. Therefore, the composition of the intestinal microbiota potentially links with the development of allergic diseases^9–11^.

In the present cross-sectional study, we investigated the relationship between daily nutritional intake, gut microbiome composition, and self-reported AR symptoms in middle-aged workers. Our aim was three-fold: (i) to suggest which nutrients are associated with AR; (ii) to search for microbes whose abundances are associated with AR; and (iii) to examine whether nutrients and microbes have a combinatorial effect on AR. Various dietary components are metabolised by gut microbiome enzymes; and therefore, the effect of dietary nutrients on the odds of AR may be modulated by intestinal microbiota.

## Methods

### Study subjects

The study participants were recruited at the Hitachi Health Care (HHC) Center, where annual health examinations of employees and their spouses from 35 affiliated companies (~38,000 workers in total) are performed. Most visitors at the HHC Center were employed by electronics-related companies and performed generally representative (not industry-specific) jobs, including general affairs, design, research and development, sales, and manufacturing. The characteristics of the visitors were described previously^12–14^. They have continuously undergone annual comprehensive health examinations, including cardiovascular and cancer screening. Therefore, we could select visitors based on the results of their previous health examinations. Previous visitors without a history of serious illness (cancer, cerebrovascular disease, or myocardial infarction) and unhealthy risk factors (such as high blood pressure, high glucose level, or abnormal lipid levels) were invited to participate in this study by e-mail. The number of enrolled volunteers was 301, and they were asked at visit reservation whether they had symptoms of seasonal or perennial AR. They re-visited the HHC Center for their annual health check-up from September 2018 to February 2019. The study participants were asked to complete a questionnaire on their dietary habits and symptoms of AR. In addition, stool samples were collected. Nine subjects were excluded due to withdrawal of consent (*n* = 2) or their inconsistent report of AR symptoms in the questionnaire compared to that at visit reservation (*n* = 7). Finally, 186 participants with symptoms of AR and 106 control subjects without symptoms of AR were included in the study. This study was reviewed and approved in advance by the Hitachi Hospital Group Ethics Committee (Approved No. 2018-5, 2019-10, and 2020-88), the Institutional Review Board of the Hitachi Ltd. (Approved No. 220-1 and 238-1), and the Research Ethics Committee of RIKEN (Approved No. H30-5). All participants provided written informed consent. This study was conducted according to the principles of the Declaration of Helsinki.

### Nutritional intake

Nutritional consumption data for the study participants were obtained using the brief-type self-administered diet history questionnaire (BDHQ). The questionnaire asked about the intake frequency of 58 selected food and beverage items, which are commonly consumed in Japan, in the preceding month^15,16^. The BDHQ does not consider the portion size. Instead, standard portion sizes were defined according to recipe books for Japanese diets^15,16^. Based on the standard tables of food composition in Japan revised in 2010^17^, crude estimates for dietary intake of total energy and 42 selected nutrients were calculated. Energy adjustment was performed using the density method^16,18^. Adjusted values for energy-providing nutrients were calculated by dividing the amount of energy obtained from each nutrient by the total energy intake. Adjusted values for non-energy-providing nutrients were defined as the amount of weight of each nutrient per 10 MJ of dietary energy intake.

### Gut microbiome

Fresh faecal samples were collected in plastic containers containing glass beads (Tomy Seiko) and RNAlater Reagent (Life Technologies Japan). The samples were transported at 4 °C to the laboratory. In the laboratory, the faecal samples (~0.2 g) were suspended in 15 mL phosphate-buffered saline (PBS) buffer and filtered with a 100-μm-mesh nylon filter (Corning) to remove human and eukaryotic cells and debris from the faecal sample. The debris on the filter was washed twice with PBS. The bacteria-enriched pellet was obtained by centrifugation of the filtrate at 9,000 × *g* for 10 min at 4 °C. The pellets were washed with 35 mL PBS once, further washed with TE20 buffer (10 mM Tris-HCl, 20 mM EDTA), and subjected to DNA extraction. Bacterial DNA was isolated and purified from the faecal samples according to enzymatic lysis methods described previously^19^.

The 16S V1–V2 region was amplified using polymerase chain reaction (PCR) with barcoded 27Fmod (5’-agrgtttgatymtggctcag-3’) and the reverse primer 338R (5’-tgctgcctcccgtaggagt-3’)^19^. PCR was performed using 50 μL of 1× Ex Taq PCR buffer composed of 10 mM Tris.HCl (pH 8.3), 50 mM KCl, and 1.5 mM MgCl_2_ in the presence of 250 μM dNTPs, 1 unit Ex Taq polymerase (Takara Bio), forward and reverse primers (0.2 μM), and ~20 ng of template DNA. PCR was performed in a 9700 PCR System (Life Technologies Japan), and the following cycling conditions were used: initial denaturation at 96 °C for 2 min, followed by 20 cycles of denaturation at 96 °C for 30 s, annealing at 55 °C for 45 s, and extension at 72 °C for 1 min, with a final extension at 72 °C. PCR amplicons were purified using AMPure XP magnetic purification beads (Beckman Coulter) and quantified using the Quant-iT PicoGreen dsDNA Assay Kit (Life Technologies Japan). An equal amount of each PCR amplicon was mixed and subjected to multiplexed amplicon sequencing with MiSeq (2 × 300 paired-end run), according to the manufacturer’s instructions.

Two paired-end reads were merged using the fastq-join program based on overlapping sequences. Reads with an average quality value of <25 and inexact matches to both universal primers were filtered out. The reads lacking both forward and reverse primer sequences were removed. Filter-passed reads were used after trimming off both primer sequences. Reads having BLAST match lengths <90% with the representative sequence in the 16S databases (described below) were considered as chimeras and removed. Finally, filter-passed reads were used for further analysis. The 16S database was constructed from three publicly available databases: Ribosomal Database Project (RDP) (Release 11, Update 5), CORE (October 13, 2017 updated; http://microbiome.osu.edu/), and a reference genome sequence database obtained from the NCBI FTP site (ftp://ftp.ncbi.nih.gov/genbank/, April 2013). From the filter-passed reads, 10,000 high-quality reads per sample were randomly chosen. The total reads (the number of samples × 10,000) were then sorted by frequency of redundant sequences and grouped into operational taxonomic units (OTUs) using UCLUST with a sequence identity threshold of 97%. The representative sequences of the generated OTUs were subjected to a homology search against the databases mentioned above using the GLSEARCH program for taxonomic assignments. For assignment at the phylum, genus, and species levels, sequence similarity thresholds of 70%, 94%, and 97%, respectively, were applied.

Three measures of alpha-diversity (Chao 1, ACE, and Shannon’s index)^20^ were calculated based on the OTU-level bacterial composition. The genus-level relative abundance was represented as a percentage. Genera with a mean relative abundance of ≥0.2% were considered for statistical analyses.

### Statistical analysis

Differences in clinical characteristics between the AR and control groups were assessed using the Wilcoxon rank sum test for continuous variables or Fisher’s exact test for discrete variables. Similarly, potential associations of nutritional and microbial variables with AR were tested using the Wilcoxon rank sum test. For nutrients and microbial genera associated with AR, age- and sex-adjusted logistic regression analysis was performed. In the logistic regression analysis, nutritional and microbial variables were classified into four categories based on quartiles (Q1–Q4). The same definition of variable categories was also used in multivariate analyses that incorporated age, sex, multiple nutrients, and/or multiple microbial genera. Statistical interaction was tested by adding a multiplicative interaction term in an age- and sex-adjusted logistic regression model and by using the chi-squared test for the comparison with and without the interaction term. Two-sided *P* <0.05 was considered statistically significant, and two-sided *P* <0.1 was regarded as marginally significant in all analyses. All statistical analyses were performed using the R software 3.6.1 (R Foundation for Statistical Computing).

### Data availability

The data are not available for public access because of participant privacy concerns, but are available from the corresponding author on reasonable request.

## Results

### Clinical characteristics of study participants

The number of subjects who reported the symptoms of AR was 186, and the number of control subjects was 106 (Table 1). The mean age of the subjects with AR and controls was 49.2 and 50.4, respectively. Almost 90% of the subjects were male in both groups. Subjects with AR symptoms had significantly lower blood pressure and triglyceride levels than controls. No significant differences between the groups were observed in body mass index, HbA1c, serum total cholesterol, low-density lipoprotein cholesterol, and high-density lipoprotein cholesterol.

**Table 1.** Characteristics of study participants

|  | Allergic rhinitis | Control | *P*-value |
| --- | --- | --- | --- |
| Number of subjects | 186 | 106 | – |
| Age, years | 49.2 ± 7.1 | 50.4 ± 8.2 | 0.117 |
| Male, % | 87.6 | 91.5 | 0.338 |
| Body mass index, kg/m<sup>2</sup> | 23.6 ± 3.0 | 23.7 ± 3.1 | 0.706 |
| Systolic blood pressure, mm Hg | 117.9 ± 11.4 | 122.4 ± 12.9 | 0.006 |
| Diastolic blood pressure, mm Hg | 75.5 ± 8.5 | 79.1 ± 8.9 | 0.003 |
| HbA1c, % | 5.6 ± 0.3 | 5.6 ± 0.4 | 0.241 |
| Serum total cholesterol, mmol/L | 5.43 ± 0.76 | 5.52 ± 0.74 | 0.593 |
| LDL cholesterol, mmol/L | 3.19 ± 0.68 | 3.28 ± 0.67 | 0.261 |
| HDL cholesterol, mmol/L | 1.58 ± 0.37 | 1.55 ± 0.41 | 0.251 |
| Triglyceride, mmol/L | 1.34 ± 0.96 | 1.49 ± 0.87 | 0.035 |

Values are shown as mean ± standard deviation or frequency. *P*-values were calculated using the Wilcoxon rank sum test for continuous variables or Fisher's exact test for discrete variables. HDL indicates high-density lipoprotein; LDL, low-density lipoprotein.

### Nutritional intake and AR

The total dietary energy was similar between the AR and control groups (mean of 7,819 and 7,966 kJ/day for the AR and control groups, respectively) (Table 2). Among the 42 selected nutrients, energy-adjusted intakes of 4 nutrients (retinol, vitamin A, cryptoxanthin, and copper) were significantly different between AR and control groups (*P* = 0.001, 0.003, 0.007, and 0.023, respectively; Wilcoxon rank sum test).

**Table 2.**
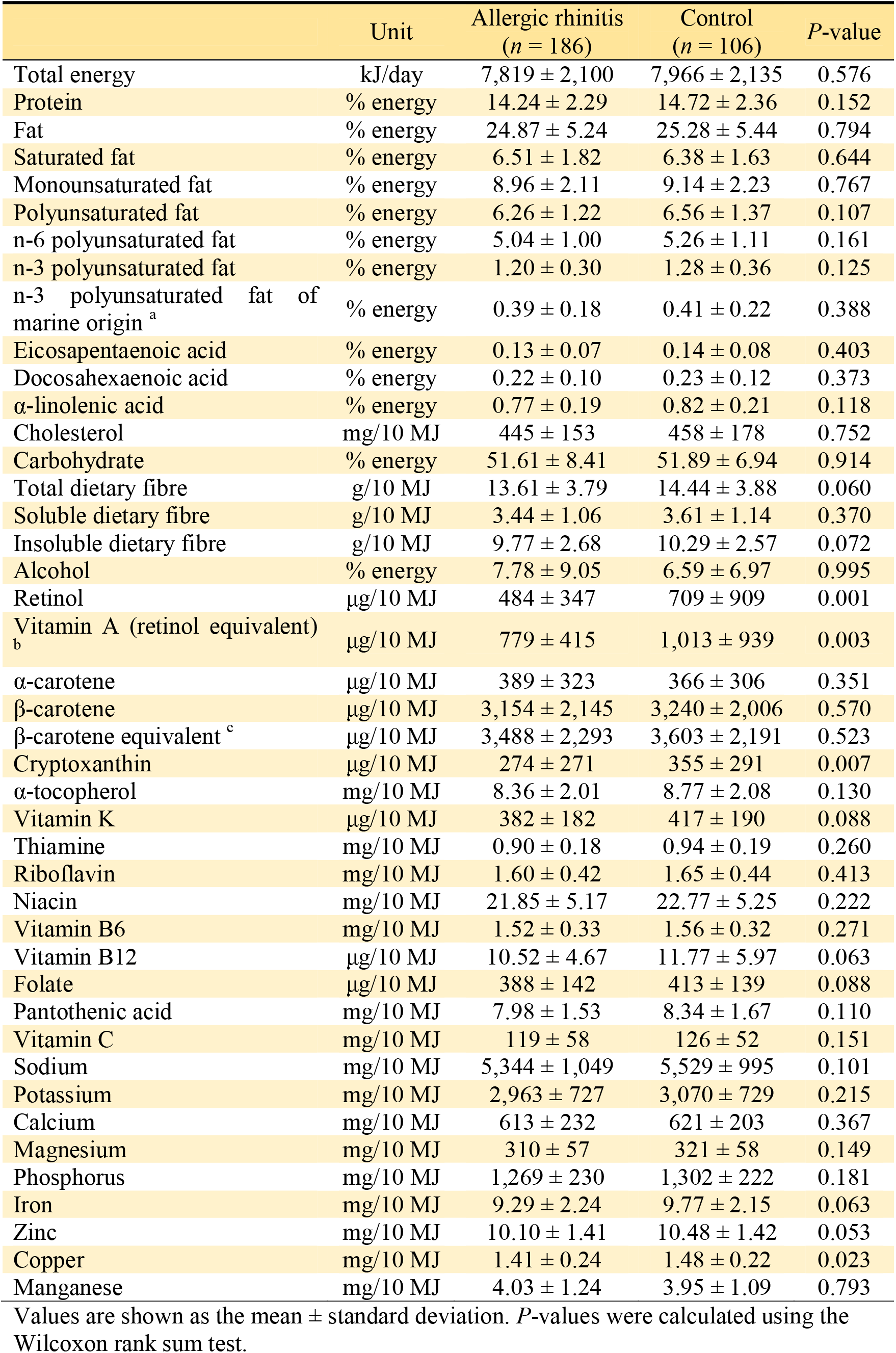
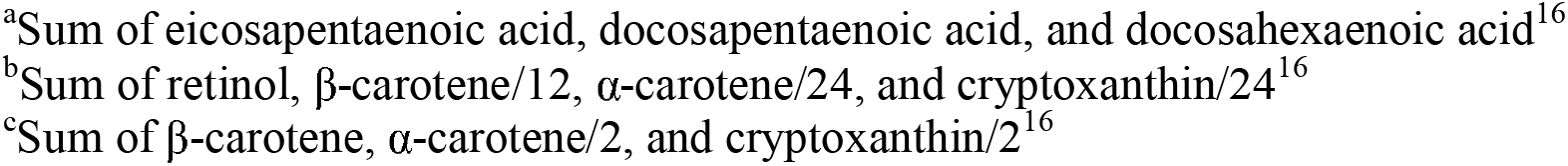
Energy-adjusted nutrient intakes and allergic rhinitis

The AR group consumed a lower level of retinol than controls (Fig. 1**a**). A clear dose-dependent trend was observed in the association between retinol and AR (Fig. 1**b**). Compared to the first quartile (Q1) of retinol intake as reference, the age- and sex-adjusted odds ratios for Q2, Q3, and Q4 were 0.83, 0.69, and 0.22, respectively (*P* for trend = 0.0004). Q4 had a significantly lower odds of AR than Q1 (*P* = 0.0001), whereas Q2 and Q3 did not. Similar dose-dependent trends were observed in the association of vitamin A and cryptoxanthin with AR (Figs. 1**c**–1**f**).

**Figure 1.**
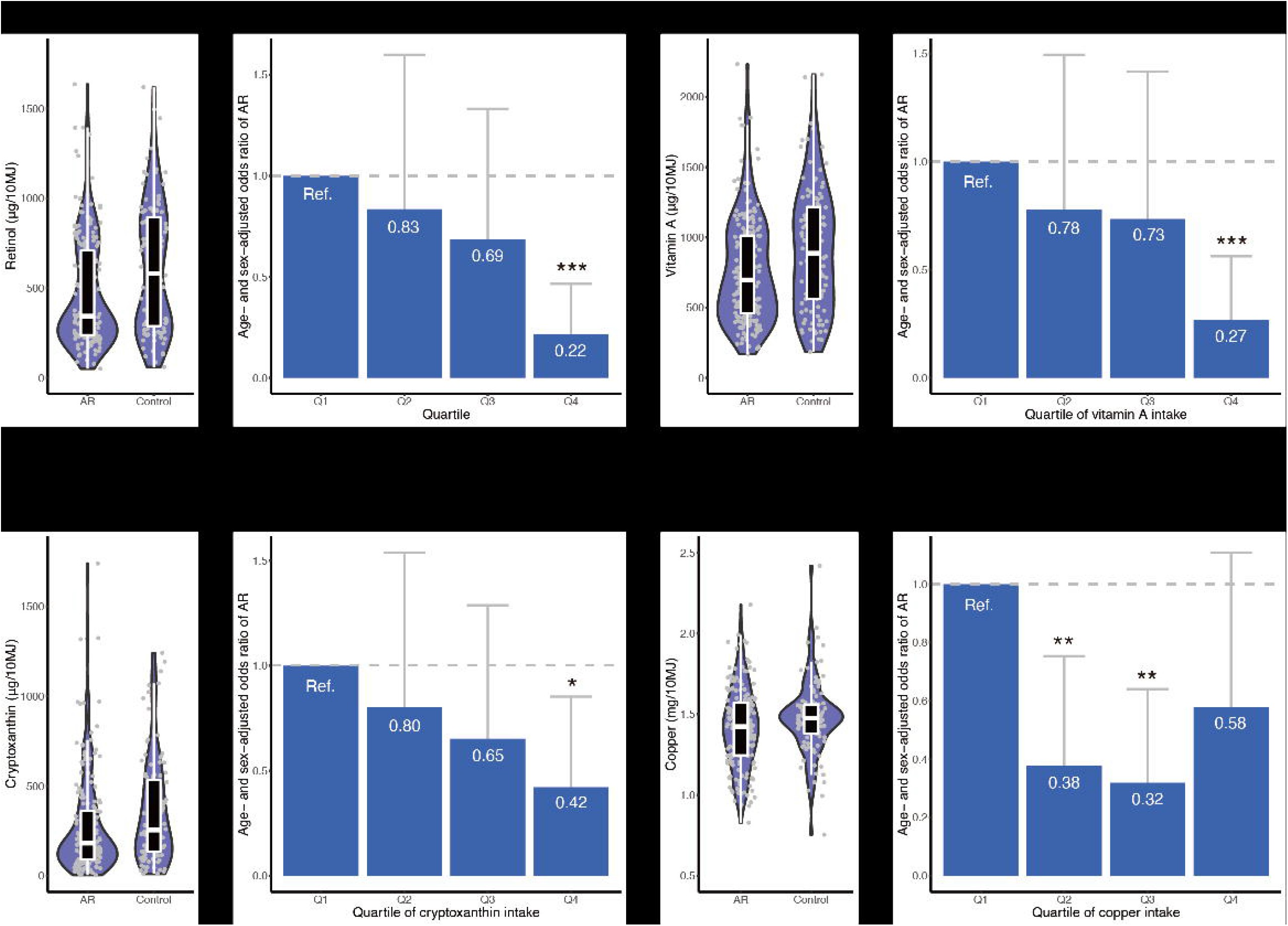
Nutrients associated with allergic rhinitis. The distribution of energy-adjusted nutritional intake is shown for (**a**) retinol, (**c**) vitamin A, (**e**) cryptoxanthin, and (**g**) copper. Outlier data points are excluded from these plots. The age- and sex-adjusted odds ratio of allergic rhinitis is presented for (**b**) retinol, (**d**) vitamin A, (**f**) cryptoxanthin, and (**h**) copper. Error bars indicate 95% confidence interval. ‘*’ indicates *P* < 0.05; ‘**’, *P* < 0.01; and ‘***’, *P* < 0.001. AR, allergic rhinitis; Q, quartile; Ref., reference group.

There was a reverse J-shaped relationship between copper and AR (Figs. 1**g** and 1**h**). Compared to Q1 of copper intake as reference, the age- and sex-adjusted odds ratios for Q2, Q3, and Q4 were 0.38, 0.32, and 0.58, respectively. Q2 and Q3 showed a significantly lower odds of AR than Q1, whereas Q4 did not.

A multivariate analysis incorporating age, sex, and the four nutrients associated with AR (*i.e.*, retinol, vitamin A, cryptoxanthin, and copper) suggested that a high level of retinol intake (Q4), a high level of cryptoxanthin (Q4), and intermediate levels of copper intake (Q2 and Q3) were independently associated with decreased odds of AR (*P* < 0.05; Supplementary Table S1). Vitamin A was not significantly associated with AR in the multivariate analysis (*P* > 0.05), possibly because the intake levels of retinol and vitamin A were highly correlated (*R* = 0.96; Supplementary Fig. S1).

### Gut microbiome and AR

Four measures of microbial alpha-diversity were not significantly different between the AR and control groups (Table 3). In total, 40 genera with mean relative abundance exceeding 0.2% were included. Among them, the relative abundances of three genera (*Prevotella*, *Bifidobacterium*, and *Escherichia*) were significantly different between the AR and control groups (*P* = 0.015, 0.048, and 0.001, respectively; Wilcoxon rank sum test).

**Table 3.** Gut microbial characteristics and allergic rhinitis

|  |  | Allergic rhinitis<br>(n = 186) | Control<br>(n = 106) | P-value |
| --- | --- | --- | --- | --- |
| Alpha-diversity | Number of OTUs observed | 295.3 ± 98.9 | 304.3 ± 100.0 | 0.412 |
|  | Chao 1 | 518.9 ± 206.0 | 548.4 ± 215.7 | 0.224 |
|  | ACE | 528.2 ± 217.2 | 557.1 ± 221.3 | 0.246 |
|  | Shannon's index | 5.04 ± 0.64 | 5.07 ± 0.55 | 0.978 |
| Genus-level<br>relative<br>abundance, % | *Bacteroides* | 19.44 ± 11.28 | 18.17 ± 11.45 | 0.440 |
|  | *Prevotella* | 6.49 ± 13.99 | 11.72 ± 17.52 | 0.015 |
|  | *Blautia* | 11.20 ± 6.14 | 10.23 ± 6.65 | 0.070 |
|  | *Bifidobacterium* | 10.41 ± 10.53 | 7.97 ± 9.57 | 0.048 |
|  | *Fusicatenibacter* | 3.85 ± 3.56 | 4.15 ± 3.95 | 0.717 |
|  | *Faecalibacterium* | 3.49 ± 2.81 | 3.66 ± 3.25 | 0.951 |
|  | *Collinsella* | 4.47 ± 5.08 | 3.30 ± 3.27 | 0.072 |
|  | *Parabacteroides* | 2.35 ± 2.19 | 3.06 ± 3.79 | 0.156 |
|  | *Megamonas* | 2.38 ± 5.87 | 2.19 ± 5.65 | 0.859 |
|  | *Streptococcus* | 2.23 ± 4.67 | 2.08 ± 4.34 | 0.224 |
|  | *Anaerostipes* | 2.19 ± 2.52 | 1.75 ± 2.01 | 0.177 |
|  | *Roseburia* | 1.73 ± 2.44 | 1.61 ± 1.73 | 0.585 |
|  | *Lachnoclostridium* | 1.39 ± 1.40 | 1.58 ± 1.73 | 0.161 |
|  | *Holdemanella* | 1.31 ± 3.67 | 1.08 ± 2.35 | 0.103 |
|  | *Dorea* | 1.03 ± 0.82 | 0.95 ± 0.87 | 0.397 |
|  | *Fusobacterium* | 0.80 ± 2.69 | 0.94 ± 1.93 | 0.113 |
|  | *Faecalicatena* | 1.07 ± 1.33 | 0.89 ± 1.19 | 0.401 |
|  | *Eubacterium* | 0.61 ± 0.67 | 0.84 ± 1.22 | 0.505 |
|  | *Catenibacterium* | 0.57 ± 2.16 | 0.72 ± 2.13 | 0.310 |
|  | *Sutterella* | 0.62 ± 0.88 | 0.65 ± 0.74 | 0.168 |
|  | *Phascolarctobacterium* | 0.52 ± 0.66 | 0.65 ± 0.89 | 0.681 |
|  | *Megasphaera* | 0.46 ± 1.12 | 0.63 ± 1.51 | 0.729 |
|  | *Prevotellamassilia* | 0.20 ± 0.96 | 0.53 ± 2.22 | 0.250 |
|  | *Coprococcus* | 0.54 ± 1.09 | 0.51 ± 0.96 | 0.784 |
|  | *Lactobacillus* | 0.66 ± 1.50 | 0.48 ± 0.84 | 0.887 |
|  | *Faecalimonas* | 0.35 ± 0.83 | 0.43 ± 1.07 | 0.376 |
|  | *Anaerobutyricum* | 0.38 ± 0.42 | 0.42 ± 0.45 | 0.592 |
|  | *Mitsuokella* | 0.26 ± 1.09 | 0.40 ± 1.35 | 0.578 |
|  | *Erysipelatoclostridium* | 0.40 ± 0.75 | 0.37 ± 0.80 | 0.202 |
|  | *Veillonella* | 0.50 ± 1.48 | 0.37 ± 1.12 | 0.815 |
|  | *Dialister* | 0.30 ± 0.72 | 0.36 ± 0.83 | 0.802 |
|  | *Butyricicoccus* | 0.36 ± 0.27 | 0.36 ± 0.30 | 0.838 |
|  | *Alistipes* | 0.35 ± 0.44 | 0.32 ± 0.39 | 0.911 |
|  | *Clostridium* | 0.43 ± 2.72 | 0.30 ± 0.93 | 0.182 |
|  | *Paraprevotella* | 0.19 ± 0.44 | 0.28 ± 0.59 | 0.127 |
|  | *Romboutsia* | 0.33 ± 0.81 | 0.28 ± 0.45 | 0.905 |
|  | *Escherichia* | 0.62 ± 2.60 | 0.26 ± 1.21 | 0.001 |
|  | *Robinsoniella* | 0.38 ± 0.97 | 0.26 ± 0.56 | 0.590 |
|  | *Lachnospira* | 0.20 ± 0.46 | 0.24 ± 0.49 | 0.871 |
|  | *Parasutterella* | 0.27 ± 0.58 | 0.18 ± 0.41 | 0.133 |

Values are shown as the mean ± standard deviation. P-values were calculated using the Wilcoxon rank sum test. OTU indicates operational taxonomic unit.

The relative abundance of *Prevotella* was either zero or near zero in a large proportion of subjects (Fig. 2**a**): 35.5% of the AR group and 25.5% of the control group did not harbour *Prevotella* at all (*i.e.*, relative abundance was 0.0%) in their gut microbial community, whereas the relative abundance of *Prevotella* widely ranged in other subjects (relative abundance reached as high as 55.6%). Compared to the first quartile (Q1) of relative abundance of *Prevotella*, the age- and sex-adjusted odds ratios for Q2, Q3, and Q4 were 0.67, 0.83, and 0.35, respectively (Fig. 2**b**). The age- and sex-adjusted odds of AR was significantly lower in Q4 than in Q1 (*P* = 0.004), but not in Q2 and Q3. A significant dose-dependent trend was observed between relative abundance of *Prevotella* and the age- and sex-adjusted odds of AR (*P* for trend = 0.017).

**Figure 2.**
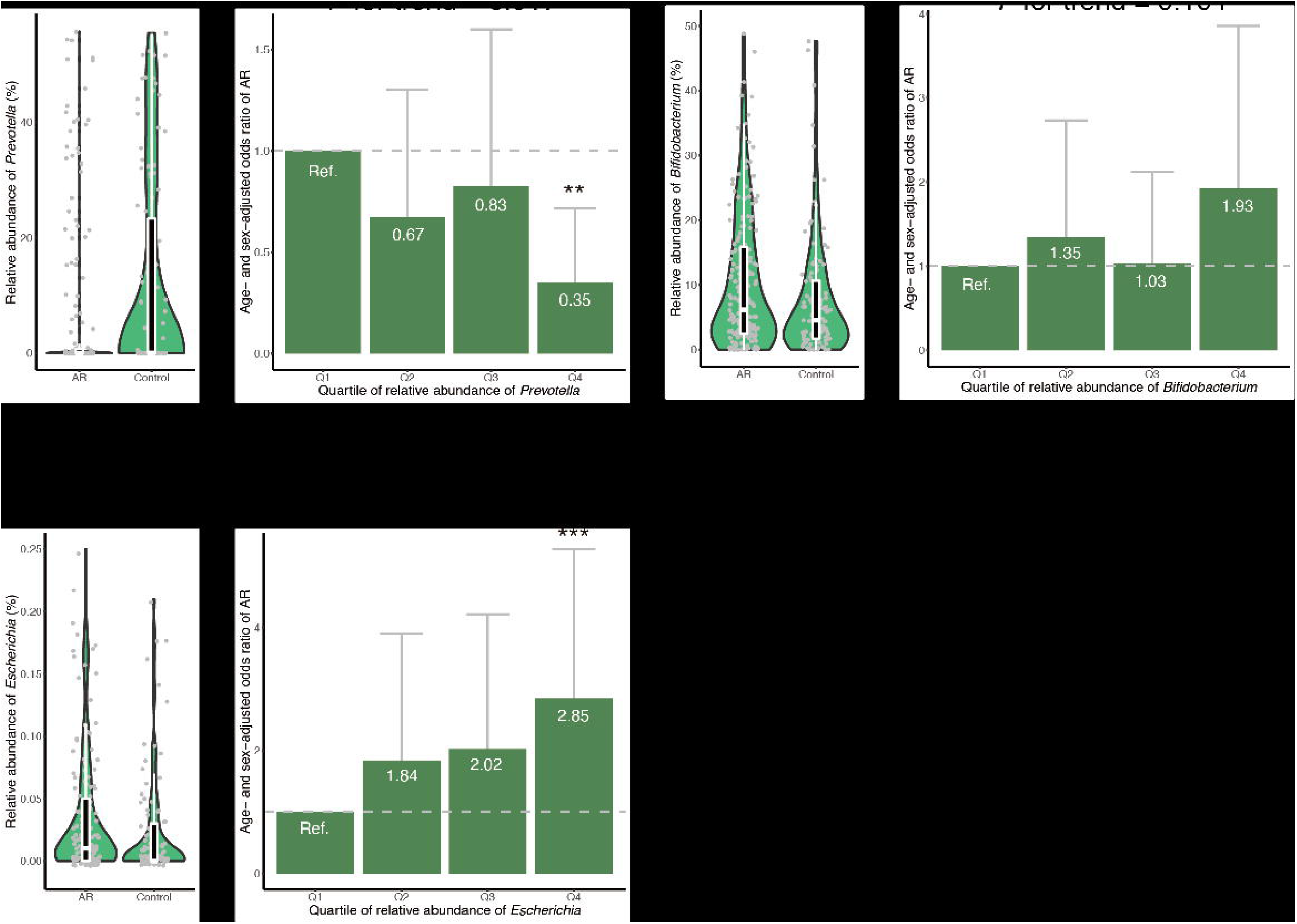
Gut microbial genera associated with allergic rhinitis. The distribution of relative abundance of gut microbial genera is shown for (**a**) *Prevotella*, (**c**) *Bifidobacterium*, and (**e**) *Escherichia*. Outlier data points are excluded from these plots. The age- and sex-adjusted odds ratio of allergic rhinitis is presented for (**b**) *Prevotella*, (**d**) *Bifidobacterium*, and (**f**) *Escherichia*. Error bars indicate 95% confidence interval. ‘*’ indicates *P* < 0.05; ‘**’, *P* < 0.01; and ‘***’, *P* < 0.001. AR, allergic rhinitis; Q, quartile; Ref., reference group.

The relative abundance of *Bifidobacterium* seemed to be higher in the AR group than in controls (Fig. 2**c**). However, dose-dependent relationship between relative abundance of *Bifidobacterium* and AR was not significant after the adjustment for age and sex (*P* for trend = 0.104). Compared to Q1 of relative abundance of *Bifidobacterium*, neither of Q2, Q3, nor Q4 showed significant difference in the age- and sex-adjusted odds of AR (Fig. 2**d**).

The AR group showed a higher relative abundance of *Escherichia* than controls (Fig. 2**e**). A clear dose-dependent relationship was observed between relative abundance of *Escherichia* and age- and sex-adjusted odds of AR (*P* for trend = 0.001; Fig. 2**f**). Compared to Q1, the age- and sex-adjusted odds ratios for Q2, Q3, and Q4 were 1.84, 2.02, and 2.85, respectively. Q4 showed a significantly higher age- and sex-adjusted odds of AR than Q1 (*P* = 0.001), whereas Q2 and Q3 did not.

No significant correlation was observed between relative abundances of *Prevotella* and *Escherichia* (*R* = −0.04; *P* = 0.477; Supplementary Fig. S2). A multivariate analysis incorporating age, sex, *Prevotella*, and *Escherichia* revealed that a high relative abundance of *Prevotella* (Q4) and a high relative abundance of *Escherichia* (Q4) were independently associated with decreased and increased odds of AR, respectively (*P* < 0.05; Supplementary Table S2). *Bifidobacterium* was not incorporated in the multivariate analysis because relative abundance of *Bifidobacterium* was not significantly associated with AR after adjusted for age and sex.

### Combinatorial effect of dietary nutrients and gut microbiome

Significant correlation was not observed between nutritional and microbial variables except for those between retinol and *Prevotella* (*R* = 0.16; *P* = 0.005) and between vitamin A and *Prevotella* (*R* = 0.13; *P* = 0.032) (Supplementary Fig. S3). A multivariate analysis incorporating age, sex, four nutrients (retinol, vitamin A, cryptoxanthin, and copper), and two microbial genera (*Prevotella* and *Escherichia*) indicated that a high level of retinol intake (Q4), intermediate levels of copper intake (Q2 and Q3), a high relative abundance of *Prevotella* (Q4), and a high relative abundance of *Escherichia* (Q4) were independently associated with AR (*P* < 0.05; Supplementary Table S3). A high level of cryptoxanthin (Q4) was marginally associated with AR in the multivariate analysis (*P* < 0.10), whereas vitamin A was not statistically associated with the AR (Supplementary Table S3).

To investigate the combinatorial effects of dietary nutrients and gut microbiome, we re-defined nutritional groups based on the abovementioned trends observed in the association analysis. Q1–Q3 were compared to Q4 for retinol, vitamin A, and cryptoxanthin, as Q4, but not Q2 and Q3, showed a significantly lower odds of AR compared to Q1 for these three nutrients. Q1 was compared to Q2–Q4 for copper because Q2 and Q3 had a significantly lower odds of AR than Q1. Similarly, Q1–Q3 were compared to Q4 for *Prevotella* and *Escherichia*.

Statistical interactions for 8 combinations of the 4 nutrients × 2 microbial genera were tested. Then, significant interactions were observed between retinol and *Prevotella* (*P* for interaction = 0.048) and between vitamin A and *Prevotella* (*P* for interaction = 0.041), but not for the other 6 combinations (Supplementary Fig. S4).

The age- and sex-adjusted odds ratio of AR was 0.64 in the low or intermediate retinol intake group (Q1–Q3) with a high relative abundance of *Prevotella* (Q4), compared to the reference group with Q1–Q3 of retinol intake and low relative abundance of *Prevotella* (Q1–Q3) (Fig. 3**a**). The age- and sex-adjusted odds ratio of AR was 0.40 in the high retinol intake group (Q4) with low or intermediate abundance level of *Prevotella* (Q1–Q3) compared to the reference. In the Q4 retinol intake group with high abundance level of *Prevotella* (Q4), a combinatorial protective effect was observed (*P* for interaction = 0.048); the age- and sex-adjusted odds ratio for this group was 0.04, which was 6.5-times smaller than the expectation based on a simple multiplication that assumes no statistical interaction (0.04 < 0.64 × 0.40 ≈ 0.26).

**Figure 3.**
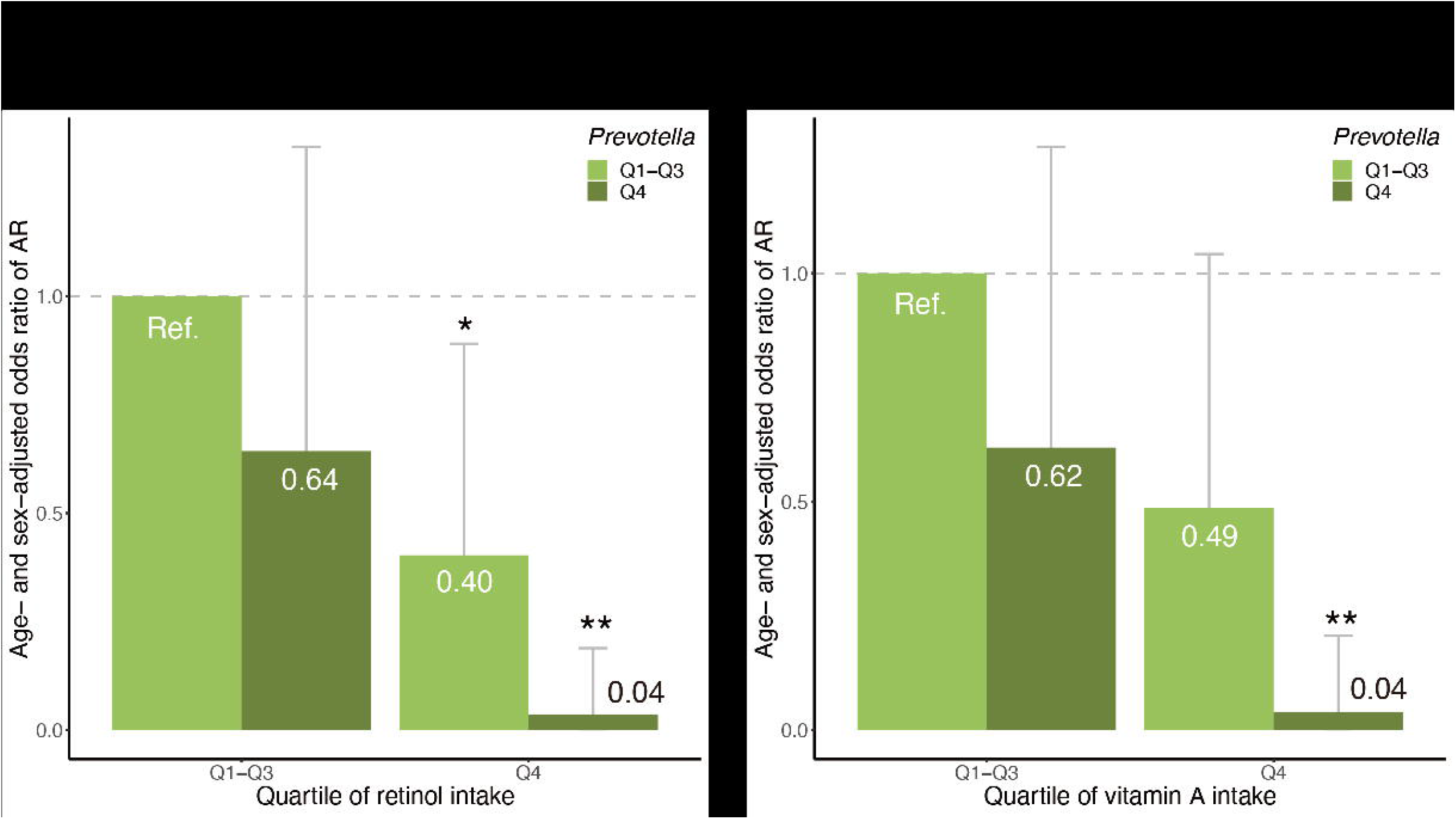
Significant statistical interactions between nutritional and gut microbial variables. Statistical interactions between (**a**) retinol and *Prevotella* and (**b**) vitamin A and *Prevotella* are shown. Error bars indicate 95% confidence interval. ‘*’ indicates *P* < 0.05; and ‘**’, *P* < 0.01. AR, allergic rhinitis; Q, quartile; Ref., reference group.

A similar statistical interaction was observed between vitamin A and *Prevotella* (Fig. 3**b**).

## Discussion

In this study, we aimed to (i) find nutrients associated with AR; (ii) discover gut microbes associated with AR; and (iii) examine the combinatorial effect between nutrients and gut microbes on AR. Towards these ends, we recruited 186 subjects with AR symptoms and 106 controls and examined their dietary nutritional intake and composition of the gut microbiome. An association analysis suggested that retinol, vitamin A, cryptoxanthin, and copper were associated with the age- and sex-adjusted odds of AR. Furthermore, the relative abundances of *Prevotella* and *Escherichia* were associated with the age- and sex-adjusted odds of AR. Lastly, statistical interactions were observed between retinol and *Prevotella* as well as between vitamin A and *Prevotella*.

Principal dietary sources of vitamin A were retinol from animal foods and provitamin A carotenoids present in fruits, vegetables, egg yolk, and butter. As in a previous study^16^, we defined dietary intake of vitamin A (retinol equivalent) as the sum of retinol, β-carotene/12, α-carotene/24, and cryptoxanthin/24. β-Carotene, α-carotene, and cryptoxanthin are common provitamin A carotenoids, which can be converted into retinol by enzymes in the intestinal mucosa^21^. As expected from this definition, dietary intake levels of vitamin A and retinol were highly correlated in our study population (*R* = 0.96). Vitamin A is essential for multiple functions in the human body, including embryonal development, good vision, epithelial differentiation, and maintenance of immune function^22^, particularly in the differentiation of naïve T cells. Without retinoic acid, a metabolite of vitamin A, transforming growth factor beta (TGF-β) promotes the differentiation of naïve T cells into T helper 17 (Th17) cells, which are involved in inflammation, autoimmunity, and allergic disorders^23,24^. In the presence of retinoic acid, TGF-β helps intestinal dendritic cells to mediate the differentiation of naïve T cells into Treg cells, which suppress autoimmune responses^23,25^. Natural Treg cells develop in the thymus, whereas Treg cells that develop in non-thymus tissues are called induced Treg (iTreg) cells^26^. iTreg cells play a crucial role in the maintenance of intestinal homeostasis, including tolerance to commensal bacteria^27^. In addition, vitamin A is important for gut mucosal turnover and barrier function^28^, intestinal IgA secretion^29^, and T cell homing to the intestine^27^. Taken together, this finding that vitamin A and retinol are associated with a decreased odds of AR may be related to the role of vitamin A in modulating intestinal immune responses.

Dietary antioxidants may protect against allergic diseases^8^. From this perspective, vitamin A and carotenoids are the most effective antioxidants at low oxygen tensions that are typical of human tissues^30^. Among the carotenoids examined in this study, only cryptoxanthin was associated with AR, whereas α-carotene and β-carotene were not. As all three of these carotenoids have antioxidant activity^31,32^, it remains unclear whether the association of retinol, vitamin A, and cryptoxanthin with AR is related to their antioxidant activity.

Copper, an essential trace metal, is a cofactor of many redox enzymes^33^. As such, it is involved in iron metabolism, antioxidant activity, neuropeptide synthesis, and immune function^34^. Copper is essential for humans but toxic at high levels; and therefore, both copper deficiency and excess can produce adverse health effects^35^. Copper deficiency influences the oxidant defence system, resulting in increased oxidative damage to lipids, DNA, and proteins^36^. Similarly, the toxic effects of copper at high concentrations are related to the generation of oxygen free radicals, and excess copper enhances lipid peroxidation and DNA damage^37^. In this study, we observed a reverse J-shaped relationship between copper and AR. This reverse J-shaped curve may be related to the anti- and pro-oxidant activities of copper at appropriate and high concentrations, respectively.

The human gut harbours hundreds of microbial species. According to Arumugam *et al.* (2011), compositions of the human gut microbiome are classified into three clusters, which are named enterotypes^38^. The first enterotype was characterised by *Bacteroides* (B-type), whereas the second and third enterotypes were dominated by *Prevotella* (P-type) and *Ruminococcus* (R-type), respectively. Later studies reduced the number of enterotype clusters into two, because the B- and P-type enterotypes were consistently identified, but the identification of R-type was dependent on clustering and modelling methods^39–41^. Note that the concept of enterotypes has been challenged by some researchers because the B- and P-type enterotypes were not clustered separately in large-scale datasets, but instead samples were distributed in a gradient across the B- and P-type groups^42^. Long-term dietary patterns are associated with enterotypes^39^. Individuals who habitually consume a Western diet high in protein and animal fat tended to have the B-type enterotype^39^. In contrast, vegetarian, or Mediterranean diets rich in fruits and vegetables were positively associated with P-type enterotype^43^. Additionally, plant-rich diets abundant in carbohydrates and fibres were associated with the P-type enterotype^41^. Based on these studies, it can be inferred that the relative abundance of *Prevotella* may reflect adherence to healthy dietary patterns. We observed a negative association between *Prevotella* and AR, and the association was still significant even after adjusted for AR-associated nutrients. Thus, it is unlikely that the association between *Prevotella* and AR is confounded by dietary patterns.

Several observational studies found that the P-type enterotype was frequently observed in rural or isolated populations, whereas the B-type enterotype was abundant in industrialised countries^44,45^. For example, a study of Chinese nomads showed that the prevalence of the P-type enterotype gradually decreased with the degree of urbanisation^46^. Given this gradient of *Prevotella* along with urbanisation, our finding of the association between *Prevotella* and AR reminds us of the microflora hypothesis of allergic diseases, which states that an unhealthy microbiota composition attributable to urbanisation or westernisation contributes to the development of allergies^47^. Several studies support this hypothesis. First, since the 1950s, industrialisation and urbanisation have accelerated around the world, and simultaneously, the prevalence of allergic diseases has increased in urban areas^48^. Second, exposures to animals, early day-care attendance, and increased number of siblings were associated with a decreased risk of allergen sensitisation^49^, indicating that early exposures to multiple types of microorganisms may facilitate development of the immune system^48^. Similarly, having older siblings was also associated with a decreased risk of AR in children^50^. Third, reduced exposure to microorganisms in early life is responsible for a shift in the balance between type 1 and type 2 helper T (Th1/Th2) cells towards the overactive Th2 arm, which stimulates IgE-mediated allergic responses^51^. Based on the microflora hypothesis, our data indicate that the abundance of *Prevotella* in adults may serve as a surrogate marker for early exposure to particular microorganisms that are essential for immune system development. Alternatively, *Prevotella* itself may be one of the essential microorganisms for the development of immune system.

Several studies have reported the relationship between *Escherichia* species and allergic disorders. The intestinal relative abundance of *Escherichia* was higher in children with asthma or rhinitis than in controls^52^. The positive association between *Escherichia* and asthma was replicated in another study^53^. In infants, a high abundance of *E. coli* was associated with an increased risk of atopic eczema^54^. To our knowledge, little is known about the molecular mechanisms underlying the association between *Escherichia* and allergies. We expanded previous findings in children by providing evidence of a positive association between *Escherichia* and AR in adults.

We observed a statistical interaction between retinol and *Prevotella* as well as between vitamin A and *Prevotella*, suggesting that a combination of high dietary intake of retinol and carotenoids with high abundance of *Prevotella* may have a protective effect on the development of AR. Retinoic acid, which is derived from retinol or carotenoids, modulates the intestinal immune system^55^. In humans, retinoic acid is irreversibly synthesised from retinal by a host enzyme, aldehyde dehydrogenase^56^, which is in turn reversibly produced from retinol by alcohol dehydrogenases or retinol dehydrogenase^56^, or from carotenoids by β-carotene-15, 15’-oxygenase 1^57^. Interestingly, some gut microorganisms encode enzymes that are potentially involved in retinal biosynthesis^58^. For example, *Prevotella marshii* DSM 16973 harbours a gene belonging to the *brp*/*blh* family^59^, which encodes an enzyme that produces retinal from β-carotene^60^. The observed statistical interaction of retinol and vitamin A with *Prevotella* may indicate a complex interplay of host and bacterial genes in the metabolism of retinol and carotenoids.

The present study has several limitations. First, we searched for nutritional and gut microbial factors associated with AR and incorporated 42 nutrients and 40 genera in association analyses. We did not apply multiple testing correction. Therefore, our findings should be carefully interpreted. Although we found multiple candidate factors associated with AR, further studies are needed to confirm the associations. Second, our definition of AR was based on self-reported symptoms. A clinical diagnosis based on interviews, rhinoscopy, skin tests, and allergen-specific IgE tests is required for a more accurate definition. Third, ~90% of the subjects were male. Further research is warranted to reveal whether our findings are consistent with female populations.

In conclusion, we suggest that four nutrients (retinol, vitamin A, cryptoxanthin, and copper) and two gut microbial genera (*Prevotella* and *Escherichia*) were associated with the age- and sex-adjusted odds of AR. In addition, a combinatorial protective effect of retinol and *Prevotella* was observed, and the age- and sex-adjusted odds of AR was 25-fold lower in subjects with a high level of dietary retinol intake and a high abundance of *Prevotella* compared to those with low retinol intake and a low abundance of *Prevotella*. Our results provide insight into the complex interplay between dietary nutrients, gut microbiome, intestinal immune systems, and the development of AR.

## Acknowledgements

The authors thank the Mr. Mitsunori Yasugi in the Hitachi Health Care Center, Hitachi Ltd. for his cooperation in collecting the faecal samples and clinical data from the study participants. This research did not receive any specific grant from funding agencies in either the public, commercial, or not-for-profit sectors.

## Author Contributions

Y.S., M.S., W.S., and T.H. wrote the manuscript. T.N. oversaw recruitment and clinical data management. Y.S., F.H., and T.H. managed the nutritional data. W.S. and Y.O. performed library preparation, sequencing, and bioinformatic analysis to obtain gut microbiome data. Y.S. and T.H. performed the statistical analysis. M.S., E.K., and M.H. supervised the work. Y.S., F.H., M.S., T.N., and M.H. designed and coordinated the project. All authors commented on and approved the manuscript.

