## Supplementary information for "Multiple nutritional and gut microbial factors associated with allergic rhinitis: the Hitachi Health Study"

Supplementary Figures S1 to S4

Supplementary Tables S1 to S3

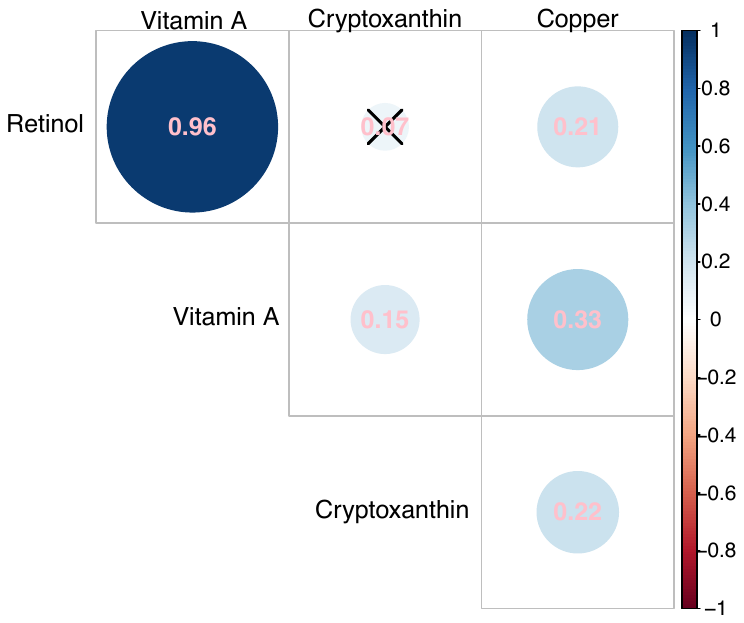

**Supplementary Figure S1.** Correlation between nutritional variables. The colour and size of circles indicate the strength of correlation between nutritional variables. Pearson correlation coefficients are shown in the circles. ‘**×**’ in the circle indicates a non-significant correlation (*P* < 0.05).

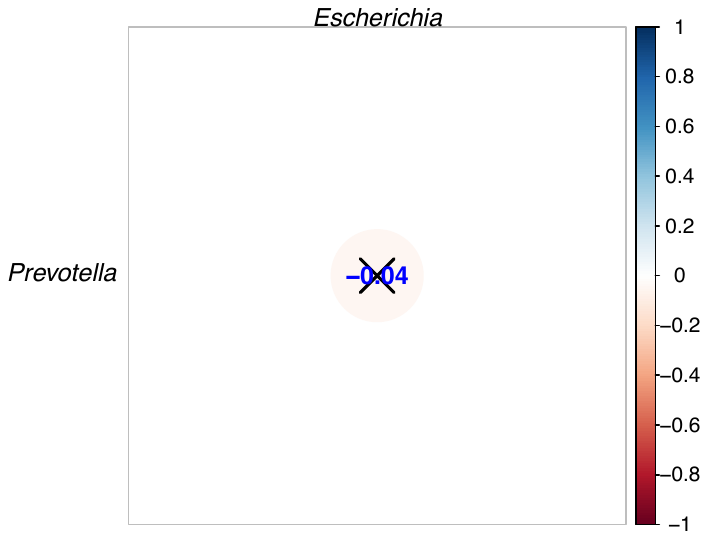

**Supplementary Figure S2.** Correlation between gut microbial variables. The colour and size of circles indicate the strength of correlation between nutritional variables. Pearson correlation coefficients are shown in the circles. ‘**×**’ in the circle indicates a non-significant correlation (*P* < 0.05).

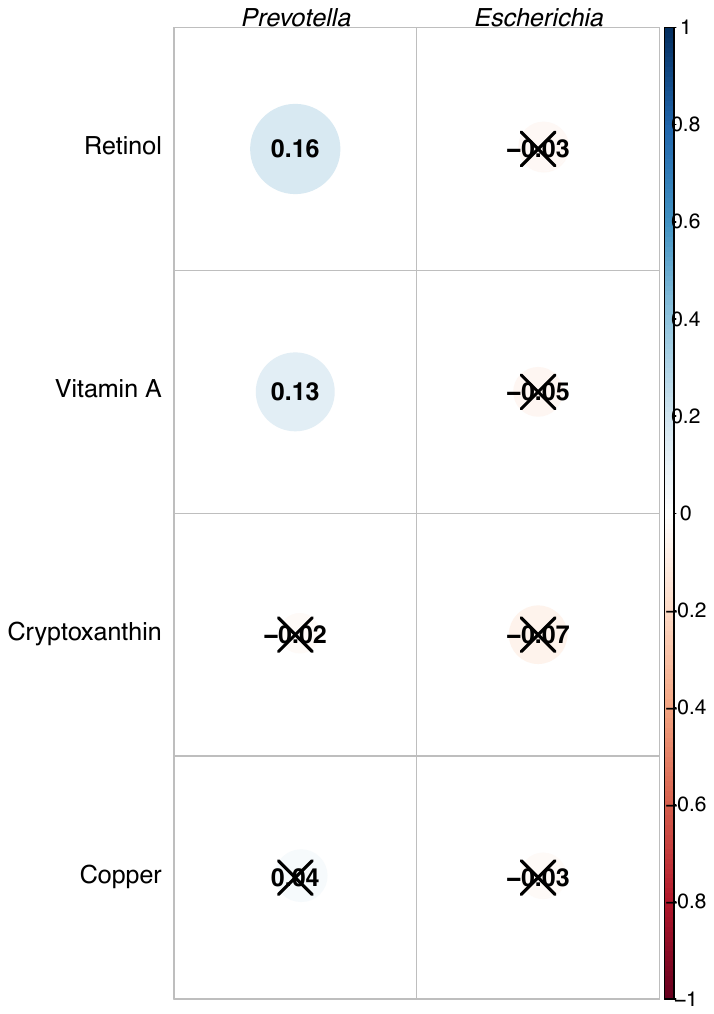

**Supplementary Figure S3.** Correlation between nutritional and gut microbial variables. The colour and size of circles indicate the strength of correlation between nutritional and microbial variables. Pearson correlation coefficients are shown in the circles. ‘**×**’ in the circle indicates a non-significant correlation (*P* < 0.05).

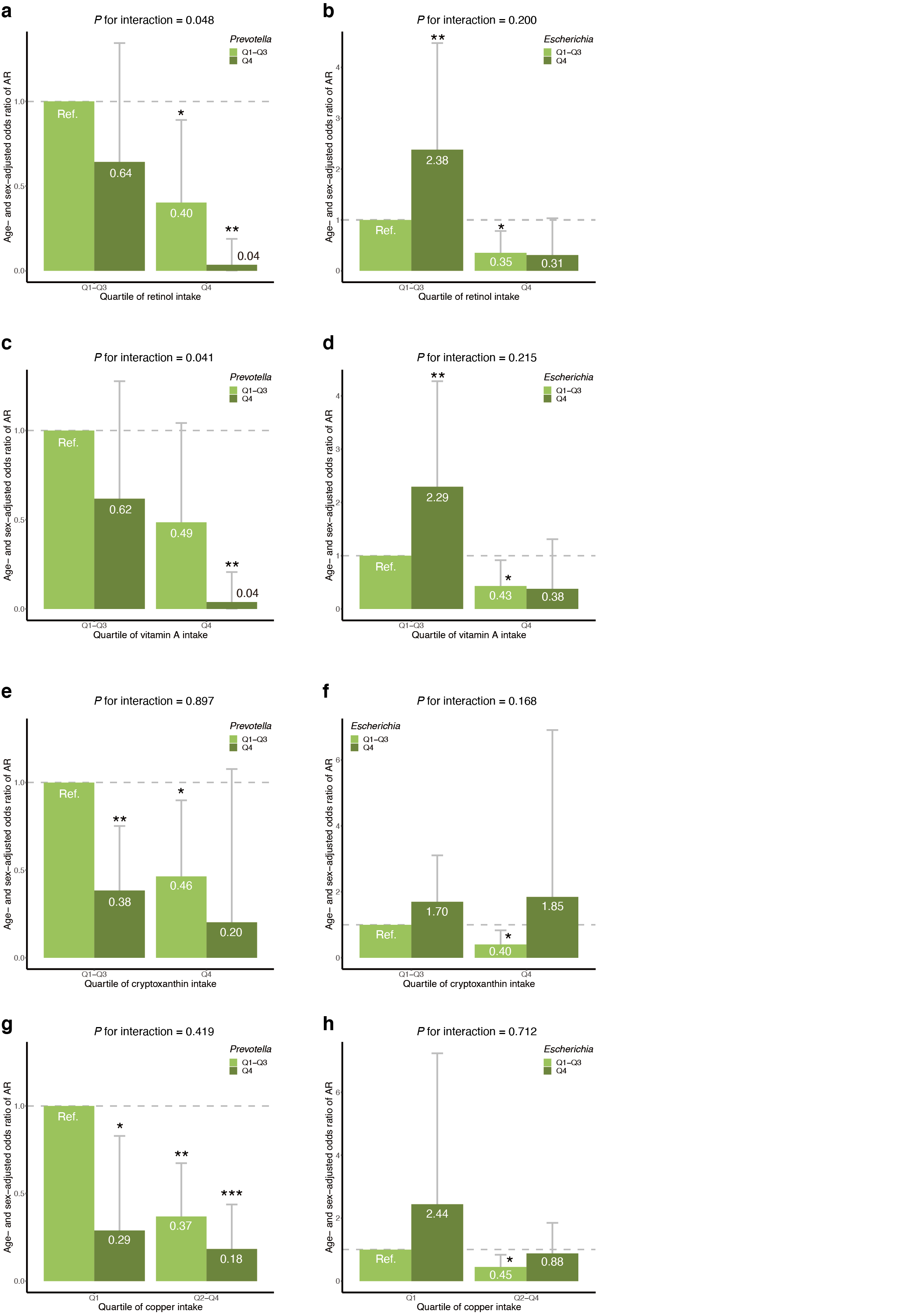

**Supplementary Figure S4.** Statistical interactions between nutritional and gut microbial variables. Statistical interaction is shown for (**a**) retinol and *Prevotella*, (**b**) retinol and *Escherichia,* (**c**) vitamin A and *Prevotella,* (**d**) vitamin A and *Escherichia,* (**e**) cryptoxanthin and *Prevotella,* (**f**) cryptoxanthin and *Escherichia*, (**g**) copper and *Prevotella,* and (**h**) copper and *Escherichia*. Error bars indicate the 95% confidence interval. ‘*’ indicates *P* < 0.05; ‘**’, *P* < 0.01; and ‘***’, *P* < 0.001. AR, allergic rhinitis; Q, quartile; Ref., reference group.

**Supplementary Table S1. A multivariate analysis of nutritional variables**

| Variable | Group | OR (95% CI) | *P* | *P* for trend |
| --- | --- | --- | --- | --- |
| Retinol | Q1 | Reference | – | 0.013 |
|  | Q2 | 0.65 (0.30–1.41) | 0.277 |  |
|  | Q3 | 0.49 (0.17–1.38) | 0.182 |  |
|  | Q4 | 0.14 (0.03–0.54) | 0.006 |  |
| Vitamin A | Q1 | Reference | – | 0.338 |
|  | Q2 | 1.40 (0.62–3.20) | 0.425 |  |
|  | Q3 | 1.85 (0.64–5.50) | 0.257 |  |
|  | Q4 | 1.74 (0.43–7.56) | 0.447 |  |
| Cryptoxanthin | Q1 | Reference | – | 0.056 |
|  | Q2 | 0.98 (0.49–2.00) | 0.964 |  |
|  | Q3 | 0.80 (0.38–1.73) | 0.575 |  |
|  | Q4 | 0.45 (0.20–0.99) | 0.048 |  |
| Copper | Q1 | Reference | – | 0.168 |
|  | Q2 | 0.35 (0.17–0.74) | 0.006 |  |
|  | Q3 | 0.31 (0.14–0.65) | 0.002 |  |
|  | Q4 | 0.68 (0.33–1.40) | 0.293 |  |

CI indicates confidence interval; OR, odds ratio; Q, quartile.

**Supplementary Table S2. A multivariate analysis of gut microbial variables**

| Variable | Group | OR (95% CI) | *P* | *P* for trend |
| --- | --- | --- | --- | --- |
| *Prevotella* | Q1 | Reference | – | 0.059 |
|  | Q2 | 0.70 (0.36–1.37) | 0.296 |  |
|  | Q3 | 0.92 (0.47–1.81) | 0.810 |  |
|  | Q4 | 0.41 (0.19–0.85) | 0.017 |  |
| *Escherichia* | Q1 | Reference | – | 0.002 |
|  | Q2 | 1.76 (0.84–3.78) | 0.137 |  |
|  | Q3 | 2.03 (0.99–4.28) | 0.057 |  |
|  | Q4 | 2.62 (1.42–4.91) | 0.002 |  |

CI indicates confidence interval; OR, odds ratio; Q, quartile.

**Supplementary Table S3. A multivariate analysis incorporating nutritional and gut microbial variables**

| Variable | Group | OR (95% CI) | *P* | *P* for trend |
| --- | --- | --- | --- | --- |
| Retinol | Q1 | Reference | – | 0.055 |
|  | Q2 | 0.76 (0.34–1.69) | 0.498 |  |
|  | Q3 | 0.62 (0.21–1.80) | 0.382 |  |
|  | Q4 | 0.19 (0.04–0.79) | 0.025 |  |
| Vitamin A | Q1 | Reference | – | 0.561 |
|  | Q2 | 1.24 (0.53–2.91) | 0.624 |  |
|  | Q3 | 1.40 (0.47–4.28) | 0.554 |  |
|  | Q4 | 1.47 (0.35–6.54) | 0.602 |  |
| Cryptoxanthin | Q1 | Reference | – | 0.071 |
|  | Q2 | 0.97 (0.47–2.01) | 0.937 |  |
|  | Q3 | 0.85 (0.39–1.89) | 0.686 |  |
|  | Q4 | 0.45 (0.19–1.01) | 0.056 |  |
| Copper | Q1 | Reference | – | 0.175 |
|  | Q2 | 0.35 (0.16–0.76) | 0.008 |  |
|  | Q3 | 0.30 (0.14–0.65) | 0.002 |  |
|  | Q4 | 0.69 (0.33–1.46) | 0.329 |  |
| *Prevotella* | Q1 | Reference | – | 0.109 |
|  | Q2 | 0.92 (0.44–1.92) | 0.822 |  |
|  | Q3 | 1.03 (0.50–2.12) | 0.941 |  |
|  | Q4 | 0.43 (0.19–0.96) | 0.040 |  |
| *Escherichia* | Q1 | Reference | – | 0.016 |
|  | Q2 | 1.45 (0.66–3.26) | 0.363 |  |
|  | Q3 | 1.74 (0.81–3.82) | 0.160 |  |
|  | Q4 | 2.23 (1.15–4.37) | 0.018 |  |

CI indicates confidence interval; OR, odds ratio; Q, quartile.
